# Risky Choices After Frontal Brain Injury: Differential Effects in Self vs Other-Decision Contexts

**DOI:** 10.1101/2025.11.21.689684

**Authors:** Farzad Rostami, Sarvenaz Soltani, Jordan Grafman, Raheleh Heyrani, Aryan Yazdanpanah, Amin Jahanbakhshi, Abdol-Hossein Vahabie, Seyed Vahid Shariat, AmirHussein Abdolalizadeh, Fatemeh Sadat Mirfazeli

## Abstract

Frontal lobe integrity is crucial for assessing risk and making informed decisions. This study investigated how frontal lobe lesions affect the computational mechanisms underlying risky choice, particularly when decisions impact oneself versus another person. A Patient Group of 20 individuals with frontal cortex damage and a Control Group of 20 matched individuals performed a gambling task, making accept/reject decisions on mixed-outcome gambles for themselves (“Self”) or an anonymous other (“Other”). We provide a mechanistic account of choice behavior using Prospect Theory, the leading behavioral model of decision-making under risk, to quantify parameters for utility curvature, loss aversion, and probability weighting. Behaviorally, the Patient Group accepted significantly more disadvantageous gambles for themselves than did the Control Group yet showed a trend toward greater caution when choosing for others. Prospect Theory modeling revealed a specific computational phenotype for this behavior. Compared to the Control Group, the Patient Group exhibited significantly more pronounced utility curvature (lower α, β) and more linear, less distorted probability weighting (higher γ). While patients also showed a trend toward lower loss aversion (λ), this difference was not statistically significant. This combination of altered utility and probability processing explains their paradoxical risk-seeking. These findings suggest that frontal cortex damage disrupts the computation of subjective value, leading to a distinctive decision-making profile marked by altered utility curvature and reduced sensitivity to outcome magnitudes. This computational characterization deepens our understanding of frontal lobe contributions to decision-making and can inform targeted rehabilitation strategies.

## Introduction

Risky decision-making is a cornerstone of human cognition, influencing choices from personal finances to leadership in social dilemmas. These decisions can be **self-focused**, affect only the decision-maker, or **involve others**, where outcomes impact people beyond oneself. This self–other distinction introduces unique complexity because it requires balancing self-interest with other-regarding values through distinct cognitive and neural mechanisms (Jung et al., 2013; Janowski et al., 2013). Behaviorally, people often make different risk choices for others than they do for themselves. A growing literature demonstrates systematic **self–other discrepancies** in risk preferences. For instance, one study found that individuals tend to be more risk-averse and loss-averse when choosing on behalf of someone else compared to choosing for themselves (Fareri et al., 2022). In that study, choices for others (especially close others like friends) were made more cautiously and consistently, whereas decisions for oneself allowed slightly more risk-taking. Social distance modulated this effect: when deciding for a friend (a socially closer other), participants’ choices became more risk-neutral and reliable than when deciding for a stranger (Fareri et al., 2022). These results suggest that people feel a greater sense of responsibility or caution with someone else’s outcomes, adjusting their decision strategy accordingly. Importantly, differences also emerge at the level of outcome processing. **Electrophysiological evidence** indicates that the brain evaluates the results of risky choices differently depending on who is affected. Xu (2021) reported that negative feedback in a Balloon Analogue Risk Task elicited a larger P300 component (a late positive potential indexing evaluative attention) when participants were risking their own outcome, compared to when they were risking someone else’s outcome (Xu, 2021). Taken together, these findings imply that despite deciding more cautiously for others, people allocate more attention or emotional significance to losses incurred for themselves than for others. These studies also underscore that self-versus-other decisions are behaviorally distinct and recruit partially different neural mechanisms. Social-cognitive processes (mediated by dmPFC, TPJ, etc.) intertwine with valuation processes during other-regarding decisions, leading to unique patterns of risk-taking (Zhang et al., 2019). Understanding these patterns is particularly important in clinical populations with frontal lobe damage, who may struggle with both decision-making and social cognition.

Damage to the frontal lobes, whether from trauma, stroke, or degenerative disease, often yields profound changes in decision-making, especially in situations involving risk or social judgment. The frontal lobe is essential for executive functions like planning, impulse control, and considering future consequences; thus, lesions in frontal regions can produce **disinhibition** and impaired judgment. Classic neuropsychological cases have shown that injuries to the **ventromedial/orbitofrontal PFC** led to reckless financial choices and insensitivity to future outcomes, even when basic intellect is preserved (Messimeris et al., 2023). Clinically, an **orbitofrontal syndrome** may emerge, characterized by antisocial behaviors, emotional instability, and impulsivity (Chow, 2021). Patients with orbitofrontal damage often exhibit heightened risk-taking, for example, gambling large sums despite losses, alongside a blunted concern for negative consequences. Recent lesion studies confirm that the **ventromedial prefrontal cortex (vmPFC) is crucial for regulating risk behavior**: individuals with vmPFC lesions tend to bet more money on gambles and have difficulty delaying gratification, reflecting a loss of the normal loss-aversion and self-control mechanisms (Kroker et al., 2023). They may also fail to adjust their strategy after losses (a behavior sometimes called “risk chasing” or perseveration), further highlighting deficits in feedback-based learning (Kroker et al., 2023). Frontal lobe damage not only affects risk-taking but can also disrupt the social aspects of decision-making. Because the medial and orbitofrontal sectors of the frontal cortex overlap with the social cognition network, patients with frontal injuries often struggle to interpret social information and to **empathize with others’ perspectives**.

Other prefrontal regions contribute to controlling risk-taking: the **dorsal anterior cingulate cortex (dACC)** and lateral prefrontal areas (e.g., dorsolateral and inferior frontal gyrus) are engaged in cognitive control processes such as response inhibition and conflict monitoring (Clairis & Lopez-Persem, 2023). These frontal regions help individuals “think twice” before acting, promoting cautious decisions under high stakes. Damage to these control regions undermines these regulatory processes and has been linked to increased impulsivity and maladaptive risky behavior. Individuals with lesions to the dACC or lateral PFC often struggle to inhibit prepotent responses and fail to learn from negative outcomes, resulting in disinhibited or suboptimal choices. Consistent with this, experimentally silencing the dACC in animal models causes more impulsive choices and suboptimal decision-making (White et al., 2024). Emotional and interoceptive inputs are also integrated into choice behavior via the **anterior insular cortex (AIC)**. The AIC links visceral emotional states (like stress or anxiety) with decision processes, and perturbations to the AIC can shift risk preferences. For example, experimental manipulations in animals show that stress-related changes in anterior insula activity led to greater risk-seeking, highlighting this region’s importance in adapting decisions under emotional strain (Shi et al., 2023). In summary, an extended healthy frontal network, including the orbitofrontal cortex (OFC)/vmPFC for value representation, dACC/lateral PFC for impulse control, and insula for emotional context, works in concert to guide effective risky decision-making under normal conditions, and damage to this network leads to impaired risk-taking decisions.

Consequently, a patient with a frontal lobe lesion may fail to appropriately adjust their risk-taking when making decisions on behalf of another person. For example, such a patient might engage in gambles that a neurologically healthy individual would avoid in situations where someone else’s welfare is at stake (Besnard et al., 2017). In summary, frontal lobe lesions provide a unique window into the necessity of frontal circuits for adaptive decision-making. They reveal how damage to specific neural substrates (for value representation, impulse control, or mentalizing) can lead to aberrant risk preferences. However, to date, few studies have directly examined how frontal lobe damage impacts the difference between self-focused and other-focused risky decisions. Addressing this gap can advance both scientific understanding and clinical care. The present study builds on the above literature by investigating risky decision-making in a cross-sectional study of patients with frontal lobe injuries. Our sample consists of individuals with diverse frontal lesions, including orbitofrontal, frontopolar, frontotemporal, and deep frontoparietal damage, and a demographically matched healthy control group. Focusing exclusively on frontal-damage patients (and excluding those with subcortical-only lesions) allows us to isolate the role of frontal cortical networks in modulating risky choices. We employ a computational modeling approach to capture the nuanced effects of frontal damage on decision behavior. Specifically, we use frameworks from Prospect Theory to quantify each participant’s risk preferences in parametric terms (such as their degree of loss aversion and probability weighting). This modeling approach enables us to examine not only overall shifts in risk attitude due to frontal injury but also context-dependent behavioral changes (self vs. other) and the underlying computational patterns of those decisions.

By analyzing how patients with frontal lobe lesions perform in self-oriented and other-oriented risk scenarios, we aim to shed light on the specific contribution of frontal neural circuits to socially contextualized decision-making. We expect this lesion-based investigation to provide causal evidence complementing prior correlational neuroimaging findings. Understanding these dynamics has important rehabilitation implications. If frontal damage is found to selectively impair certain aspects of decision-making (for example, exaggerating risk-seeking for oneself or failing to adjust caution when others are impacted), such knowledge can guide targeted interventions. Rehabilitation programs for frontal lobe patients might incorporate decision-making training modules that explicitly address these deficits, for instance by coaching patients to pause and consider consequences for others (to counteract impulsivity and egocentric decisions) or using feedback-based exercises to improve learning from negative outcomes. Indeed, behavioral therapy for brain-injured patients often targets issues like disinhibition and impaired social judgment (Skromanis et al., 2023). The insights from this study can refine those interventions by pinpointing which cognitive mechanisms to focus on. In sum, by examining risky choice behavior through the dual lenses of social context and frontal lobe integrity, our study seeks to advance theoretical models of decision neuroscience and contribute to better clinical management of patients with frontal lobe damage. Ultimately, deeper knowledge of how the frontal lobes govern risky decisions for self and others may help affected individuals relearn safer decision strategies and improve their autonomy and social functioning after injury.

## Materials & Methods

### Participants

A total of 33 patients with brain injuries and 30 healthy control participants were initially recruited for the study. All patients had documented brain lesions involving frontal cortical regions, and all controls were neurologically healthy. Eleven patients were excluded due to lesion characteristics (9 had primarily subcortical lesions without significant frontal cortical involvement, and 2 had unclear or diffuse diagnoses), leaving **20 patients with confirmed frontal lobe lesions** (16 men, 4 women; mean age = 35.09 years, *SD* = 11.77). These frontal lesions were confirmed via clinical neuroimaging and were mainly in the orbitofrontal or ventromedial frontal regions (with some patients having dorsal frontal involvement as well). Ten control participants were excluded to better match the patient sample (7 had exceptionally high educational attainment beyond a master’s degree, and 3 showed anomalously extreme response patterns with >90% gamble acceptance, indicating potential non-compliance). This yielded a final **healthy control sample of 20** individuals (16 men, 4 women; mean age = 37.65 years, *SD* = 9.06), demographically matched to the patients on age, sex, and general education level. All participants provided informed consent. Exclusion criteria for both groups included any history of major psychiatric illness (e.g., psychosis, mania, moderate-to-severe depression, substance abuse) or additional neurological conditions (such as epilepsy or significant stroke outside the lesion area for patients, or any neurologic disorder in controls). The healthy controls were primarily hospital staff. The patients’ frontal lobe injuries resulted from traumatic brain injury (TBI) and were chronic (all patients were in a stable post-acute phase). All patients were tested at least one-month post-lesion, ensuring that acute effects had resolved and performance reflected stable, chronic impairment. No patient had widespread cognitive impairments preventing task comprehension, as verified by neuropsychological screening.

### Risky Decision-Making Task

We employed a computerized risky choice task adapted from **Edelson et al. (2018)**’s paradigm for self-versus-other decision-making. Each participant completed two blocks of trials: one block making decisions **for oneself** (“Self” condition) and another block making decisions **on behalf of another person** (“Other” condition). The order of blocks was counterbalanced across participants to control for order effects. In each block, the participant encountered a series of 48 independent gambles. On each trial, a **mixed-valence lottery** was presented, consisting of a potential monetary gain and a potential monetary loss, each paired with a specified probability of occurring. The task display showed the probability of winning versus losing (as a percentage and a graphical bar) and the corresponding gain amount (in positive **Tomans**) versus loss amount (in negative Tomans). For example, a trial might show a **20%** chance to win **+5000** Tomans (approximately $0.40 USD) and an **80%** chance to lose **–1000** Tomans (approximately $0.08 USD). The probabilities of a win were systematically varied across four levels (20%, 40%, 60%, 80%), and outcome magnitudes were drawn from a predefined set of values. Gain amounts were in the range **+1000 to +7000 Tomans** (approximately $0.08–$0.56 USD, in increments of 2000 Tomans), and loss amounts ranged from **–1000 to –5000 Tomans** (approximately $0.08–$0.40 USD, increments of 2000 Tomans), similar to the range used by Edelson and colleagues. At the time of testing, the exchange rates were approximately 12,500 Tomans per USD. This yielded a range of gamble expected values (*EV = p(win)×gain + p(lose)×loss*) from substantially negative to substantially positive.

Each trial required a **binary choice**: the participant decided to either **accept the gamble**, in which case the outcome (win or loss) would be realized with the given probabilities, or **reject the gamble**, in which case neither gain nor loss was incurred for that trial (Figure 1). Participants indicated their choice by button press, with no time limit (though reaction times were recorded). They were explicitly instructed that in the “Self” block, accepting a gamble meant they would win or lose the stated amounts, whereas in the “Other” block, their choices would affect a **hypothetical anonymous other person** instead of themselves. Importantly, participants were told that **one trial from each block would be randomly selected at the end to count “for real”**: the outcome of a randomly chosen self-decision trial would be applied to their bonus payment, and the outcome of a randomly chosen other-decision trial would hypothetically be applied to another person’s bonus. All other trials would not directly affect payment. To ensure incentive compatibility, each participant was given a starting endowment (bankroll) of 15,000 Tomans (approximately $1.20 USD) and informed that their bonus earnings could increase or decrease based on the selected gamble outcomes. They were **kept unaware of the outcomes** of each gamble during the task to prevent feedback from influencing subsequent choices. In the “Other” condition, they were told to imagine that *someone else’s* payment would be determined by their decision on the selected trial, and likewise, to imagine that another person’s decision would determine the outcome of one of *their* own trials. Participants never met the individuals for whom they made decisions (to avoid personal familiarity effects), and all decisions for others were made anonymously. These instructions emphasized the responsibility in other-regarding decisions while maintaining the hypothetical nature of the other person. The task lasted roughly 15–20 minutes per block. Performance-based compensation was computed after task completion, and all participants received at least a base compensation with the possibility of earning more based on their gamble outcomes.

**Figure 1.**
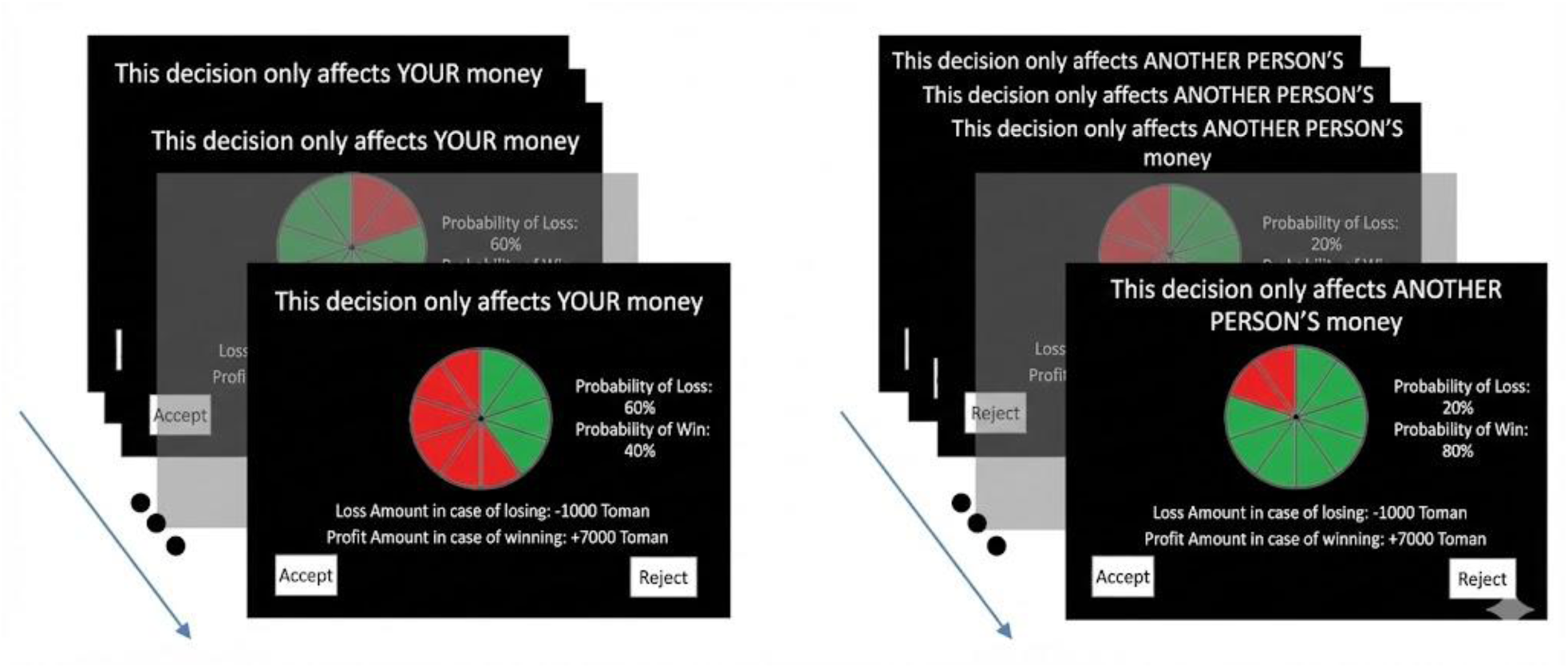
Task structure and trial presentation. (A) In the “Self” block, participants made risky decisions that directly affected their monetary outcomes. (B) In the “Other” block, participants made analogous decisions on behalf of an anonymous, unseen participant, whose payment would be determined by one randomly selected decision. Each trial presented a mixed lottery showing win/loss probabilities and magnitudes, and participants indicated whether to accept or reject the gamble. The graphical interface displayed outcome probabilities as segmented bar charts alongside textual information, ensuring clarity in value and risk representation.

### Behavioral Modeling and Analysis

#### Model-Free Behavioral Analysis

We first analyzed the risk-taking behavior descriptively by computing the acceptance rate of gambles across different conditions. For each participant, we sorted trials by the gamble’s expected value (EV) and grouped them into deciles (10 bins from most negative to most positive EV) to examine how often participants accepted gambles with negative versus positive expected value. This provided an overall behavioral risk profile, visualized as group-level acceptance curves for each decision context (Figure 2). We statistically tested for differences in mean acceptance rates using both within-subject (Self vs. Other) and between-group (Patient vs. Control) comparisons. Paired t-tests were used to compare each group’s self vs. other acceptance rates (with Bonferroni correction for multiple comparisons), and two-sample t-tests were used to compare patients vs. controls. We paid particular attention to differences in the negative-EV range, where rejecting is optimal, versus the positive-EV range, where accepting is optimal, to precisely characterize how groups differed in their choices on disadvantageous versus advantageous gambles. To formally quantify individual risk preferences and the underlying cognitive mechanisms, we applied a computational model grounded in **Prospect Theory** to each participant’s trial-by-trial choice data. This model assumes that individuals compute a subjective value for each gamble based on nonlinear transformations of utility and probability. The model includes five free parameters that capture key psychological aspects of decision-making: utility curvature for gains (α) and for losses (β), which describe the shape of the value function (α, β < 1 typically indicate diminishing sensitivity); loss aversion (λ), which scales the relative impact of losses versus gains (λ > 1 implies losses loom larger than gains); probability weighting (γ), which reflects subjective distortions of objective probabilities (γ < 1 captures the classic pattern of overweighting low probabilities and underweighting high ones); and choice consistency (θ), a softmax inverse temperature that governs the stochasticity of decisions. Under this model, the subjective value of each gamble was computed as 𝑉(𝑔𝑎𝑚𝑏𝑙𝑒) = 𝑝^𝛾^. 𝑢(𝑔𝑎𝑖𝑛) + (1 − 𝑝)^𝛾^. (−𝜆 . |𝑙𝑜𝑠𝑠|^𝛽^). The probability of accepting a given gamble was then determined by a softmax function, 𝑃(𝑎𝑐𝑐𝑒𝑝𝑡) = 1/(1 + 𝑒^(−𝜃.𝑉)^). We fit this model separately for each participant in each condition (Self and Other) by maximizing the likelihood of their trial-by-trial choices. To ensure reliable estimates, we constrained parameters to plausible ranges (e.g., 0 < α, β ≤ 2, 0 < γ ≤ 2, λ ≥ 0) and used multiple starting points for optimization. The model’s goodness of fit was confirmed by comparing its predicted acceptance rates to the observed behavior, which showed a close correspondence across all conditions (Wulff & Pachur, 2016). After fitting, we performed mixed-design ANOVAs and post-hoc t-tests on each parameter (with Bonferroni correction) to determine if frontal lesion patients differed significantly from controls in their risk attitudes and whether these parameters changed between self and other decisions, allowing us to interpret how the groups’ decision-making processes differed (Sokol-Hessner et al., 2013).

**Figure 2.**
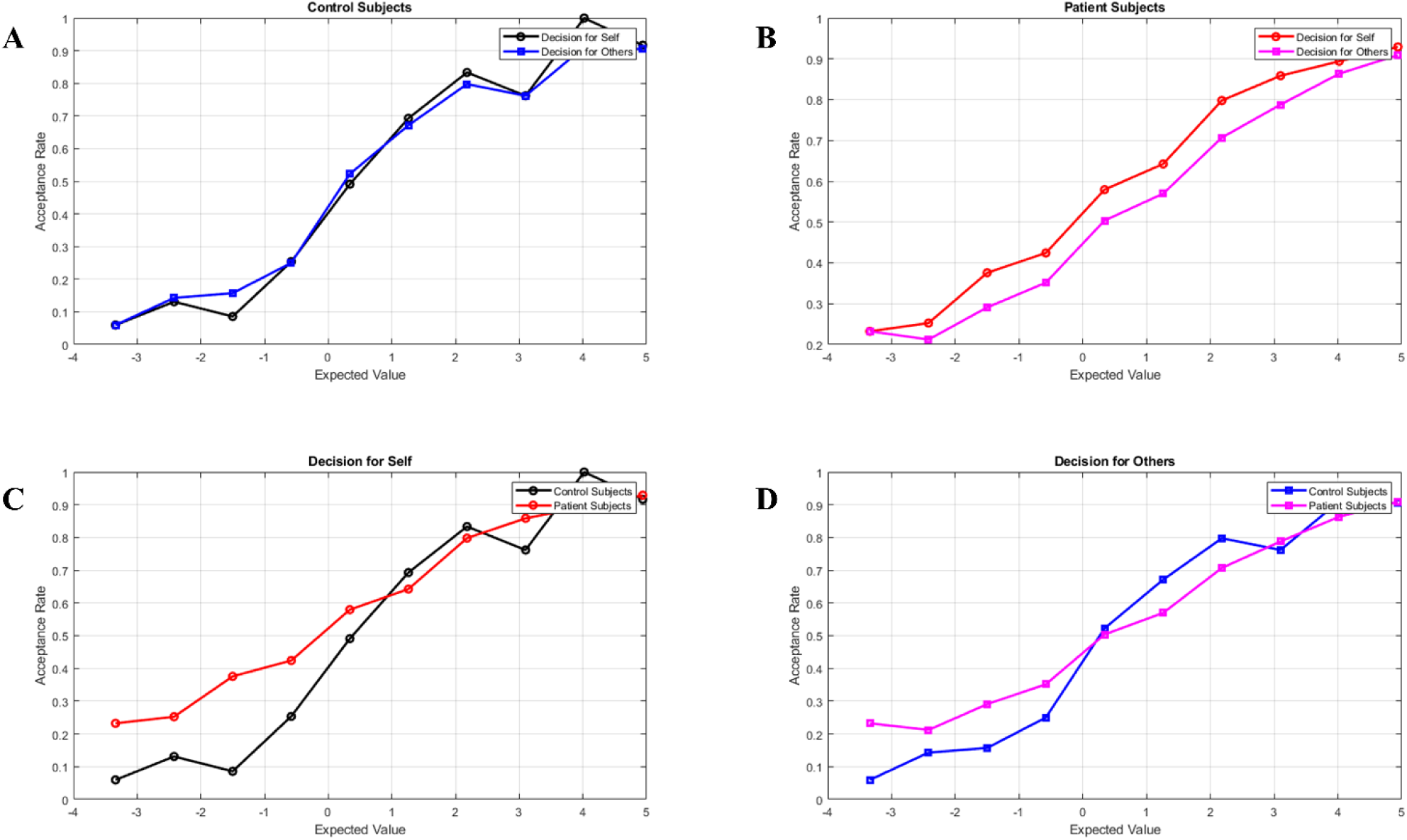
Risky decision-making behavior in patients vs. controls (Self vs Other decisions). Each panel shows the gamble acceptance rate as a function of the gamble’s expected value (EV), binned into deciles. (A) Control group – decisions for Self (black line) vs for Others (blue line). (B) Frontal lesion patient group – decisions for Self (red) vs for Others (purple). (C) Self-decision context – controls (black) vs patients (red). (D) Other-decision context – controls (blue) vs patients (purple). Healthy controls showed near-identical risk curves for Self vs Other decisions, whereas patients accepted more gambles with negative EV for themselves and slightly fewer high-EV gambles when choosing for others.

## Results

### Model-Free Behavioral Analysis

Overall, both groups’ gamble acceptance rates increased as the gambles’ expected value became more favorable, but clear group and context differences emerged (**Figure 2**). Healthy **control participants** behaved as expected for mildly risk-averse decision-makers: they rarely accepted gambles with strongly negative EVs and accepted nearly all gambles with highly positive EVs. Furthermore, controls showed no reliable difference between decisions for themselves and for another person, with paired t-tests confirming no significant difference in acceptance rates between Self and Other for either negative-EV or positive-EV gambles (all p > 0.5).

In contrast, frontal lesion patients exhibited a striking dissociation in their risk behavior. When deciding for themselves, patients were notably more willing to accept gambles with negative expected values than controls were. For these disadvantageous trials, the patient group’s acceptance rate in self-decisions was significantly higher than the control group’s (two-sample t (38) = 2.87, p = 0.0058, Cohen’s d = 0.91), indicating that frontal damage was associated with increased maladaptive risk-taking in personal decisions. Conversely, when making choices for another person, patients’ behavior shifted toward conservatism. Their acceptance curve in the Other context lay generally below their curve for Self-decisions, and although this within-subject difference did not reach significance for any single EV bin, the overall acceptance rate was marginally lower for Other decisions than for Self-decisions (paired t (19) = –2.12, p = 0.042, Cohen’s d = 0.47, uncorrected). A comprehensive three-way ANOVA on acceptance rates (Group × EV Valence × Decision Context) confirmed a significant interaction, driven by the fact that patients took more risks than controls specifically for negative-EV gambles in the Self context, but were more restrained (if anything, slightly more risk-averse than controls) in the Other context. This suggests that frontal damage disrupts not only the evaluation of personal risk but also the modulation of that risk in a social context.

### Prospect Theory Model Results

Fitting the Prospect Theory model provided insights into *which* decision-making parameters were altered by frontal lobe lesions. The model captured the data well, closely reproducing each group’s acceptance curve in all conditions (Figure 3). Despite the Patient Group’s higher propensity to choose risky options in certain situations, their **estimated utility function** parameters indicated a *more cautious* subjective valuation profile on average. Specifically, patients had significantly lower **α** and **β** parameters than the Control Group (in both Self and Other contexts). These parameters govern the curvature of the utility function for gains and losses, respectively. For gains (α), the Patient Group showed a substantially more concave utility curve than the Control Group, a statistically significant difference (mean α ≈ 0.30 vs. ∼0.65; main effect of Group: F (1, 38) = 24.83, p < 0.001, *η²* = 0.395). This indicates that patients derived diminishing additional subjective value from larger gains and were thus more risk-averse for monetary gains. For losses (β), a similar pattern emerged: the Patient Group had a lower β, implying a more convex utility curve for losses, and this group difference was also significant (mean ≈ 0.25 vs. ∼0.5; main effect of Group: F (1, 38) = 19.70, p <0.001, *η²* = 0.341). This means the Patient Group showed diminishing sensitivity to larger losses, as each additional unit of loss felt “less bad” to them compared to how it felt to controls. Taken together, the utility function of the Patient Group was more curved for both gains and losses than that of the Control Group, which would ordinarily translate to very cautious choices.

**Figure 3.**
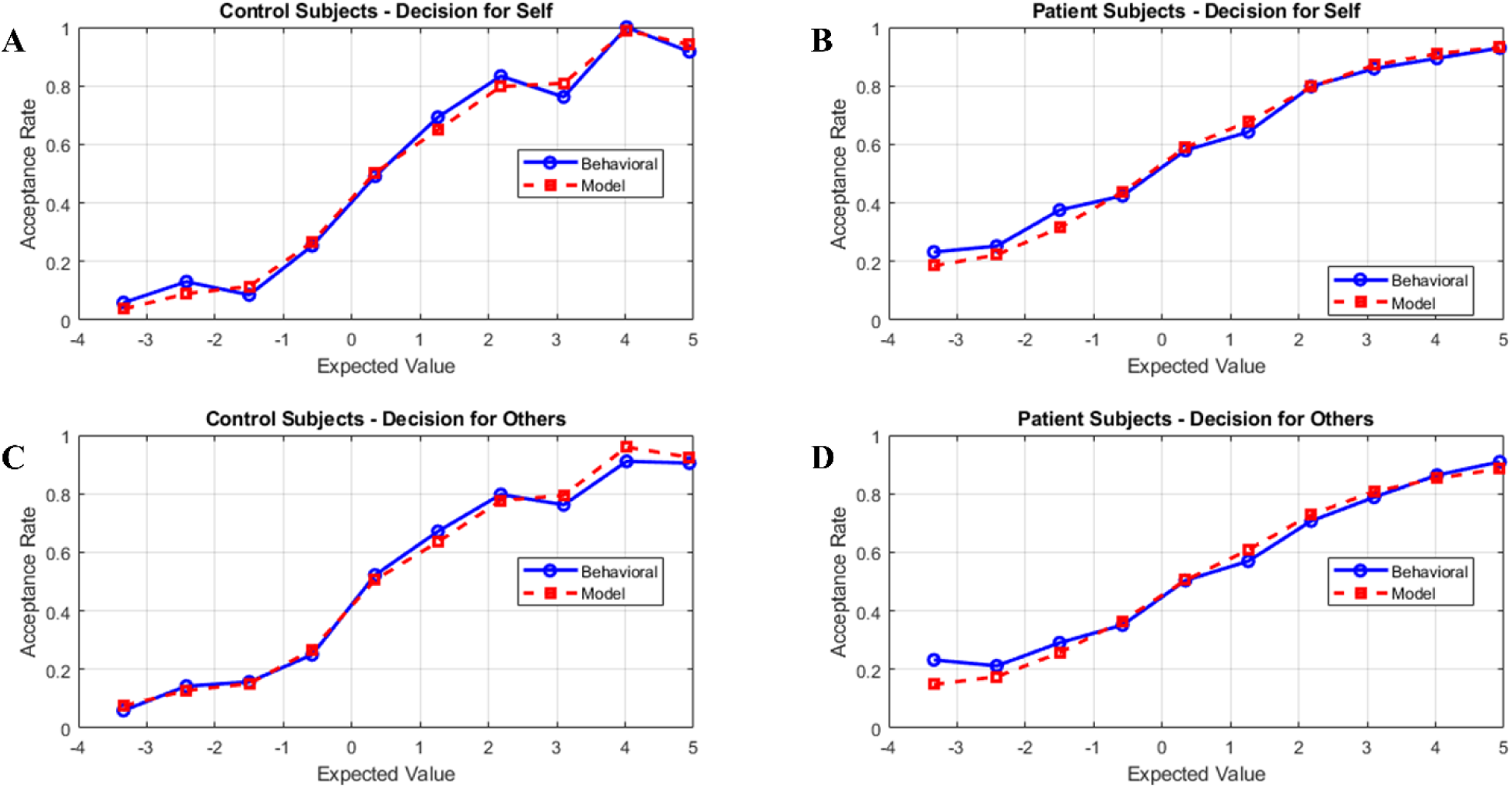
Prospect Theory model fit vs. observed behavior. Each panel compares the actual gamble acceptance rate (blue solid line, “Behavioral”) to the model-predicted acceptance rate (red dashed line, “Model”) as a function of expected value. **(A)** Control Group – Self-decisions. **(B)** Patient Group – Self-decisions. **(C)** Control Group – Other-decisions. **(D)** Patient Group – Other-decisions. In all cases, the Prospect Theory model closely approximated the participants’ choice behavior, indicating that the estimated parameters provide a good account of their risk preferences.

Despite having more curved utility functions, the **Patient Group ended up taking more risks in negative-EV scenarios**, which seems paradoxical. The model pointed to **probability weighting** as a key explanation for this paradox. The probability weighting parameter **γ** differed markedly between groups. The Control Group showed γ significantly less than 1 (around 0.30–0.40), consistent with the typical human bias to overweigh small probabilities and under-weigh large ones. The Patient Group, however, had γ values much closer to 1 (mean ≈ 0.65), and this difference was statistically significant (main effect of Group: F (1, 38) = 5.51, p = 0.022 *η²* = 0.127). This indicates that patients’ probability perception was more linear and **less biased** than controls. In practical terms, the Patient Group did not exhibit the usual probability distortions. When faced with a gamble that had, say, a 20% chance of a large gain, a control participant might subjectively overweigh this small probability, but this is often counteracted by a pessimistic distortion of the corresponding high probability of loss. The patient, by contrast, would treat the probabilities more veridically. This more linear weighting of probabilities made patients relatively less sensitive to the threat of low-probability losses. Indeed, model simulations confirmed that the increase in γ was the critical factor allowing patients to accept some gambles that controls would reject. Thus, the patients’ reduced probability distortion helped offset their generally risk-averse utility tendencies, leading to their seemingly riskier choices in negative-EV situations.

We also examined the **loss aversion** parameter **λ**. As expected, control subjects were loss-averse on average (mean λ ≈ 1.5 in the Self condition), meaning losses weighed more heavily than equal-sized gains in their decisions. While the Patient Group showed a trend toward lower loss aversion (mean λ ≈ 1.1), this difference did not reach statistical significance (main effect of Group: F (1, 38) = 2.88, p = 0.095, *η²* = 0.070). This suggests that while a reduction in the emotional impact of losses may contribute to the behavioral pattern, it was not as robust a factor as the alterations in utility curvature and probability weighting in our sample. For all parameters, there were no significant Group × Context interactions (α: F (1, 38) = 0.00, p = 0.968, *η²* < 0.001; β: F (1, 38) = 3.32, p = 0.074, *η²* = 0.080; γ: F (1, 38) = 2.23, p = 0.141, *η²* = 0.055; λ: F (1, 44) = 0.38, p = 0.617, *η²* = 0.007), indicating the group effects were fairly consistent regardless of whose outcome was at stake. Finally, the choice consistency parameter θ did not differ significantly between groups (p > 0.05 after correction), indicating that the behavioral differences stemmed from true preference shifts rather than merely random or erratic decision-making by the Patient Group.

To summarize the model findings: **frontal lobe damage caused a systematic shift in decision-making parameters**. The Patient Group showed *significantly steeper utility curvature* (lower α, β), and *more linear probability weighting* (higher γ) than controls (Figure 4). The injury affected fundamental evaluation processes across both self and other contexts, rather than simply causing indiscriminate impulsivity. In addition to the model’s good fit, we verified that the Prospect Theory model replicated the empirical choice patterns. Figure 3 shows the model’s predicted acceptance rates plotted against the actual behavioral acceptance rates for each group and condition. The model’s predictions (dashed red lines) closely tracked the observed data (solid blue lines) for both the Control Group and Patient Group, in both Self and Other contexts. This validation gives confidence that the parameter differences identified above are meaningful and not an artifact of poor model fit.

**Figure 4.**
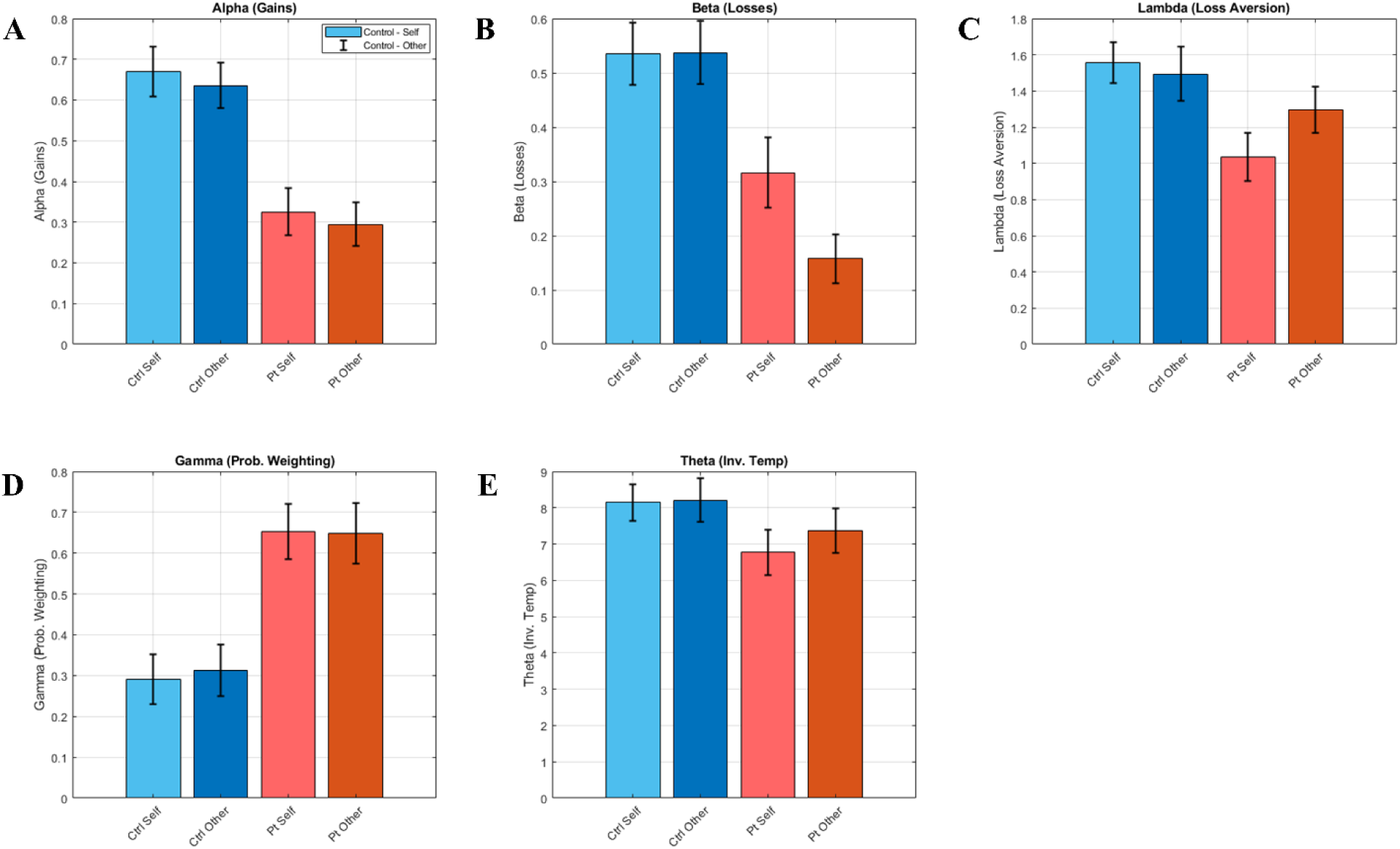
Prospect Theory model parameters for the Patient Group (red) vs. the Control Group (blue). Each plot shows the group mean (± SE) of a fitted parameter, separately for decisions affecting Self (lighter shade) and Others (darker shade). **(A) Alpha (Utility Curvature for Gains):** The Patient Group has significantly lower α. **(B) Beta (Utility Curvature for Losses):** The Patient Group’s β is lower. **(C) Lambda (Loss Aversion):** The Patient Group shows a non-significant trend toward lower λ. **(D) Gamma (Probability Weighting):** The Patient Group’s γ is higher (closer to linear weighting). **(E) Theta (Choice Consistency):** No significant group difference.

## Discussion

Our findings reveal how frontal lobe lesions fundamentally alter the cognitive and emotional processes underlying risky decision-making. Using Prospect Theory as a framework, we identified a specific computational phenotype that explains the paradoxical behavior observed in patients with frontal lobe damage. These patients exhibited significantly steeper utility curves for both gains and losses (lower α and β values), indicating a diminished sensitivity to outcome magnitudes. Concurrently, their probability weighting function became more linear (higher γ), suggesting a reduction in the typical cognitive biases that govern the perception of chance. Although patients showed a trend toward reduced loss aversion (lower λ), this was not statistically robust. Crucially, their choice consistency (θ) was intact, confirming that these behavioral differences were driven by systematically altered preferences rather than by random or noisy decision processes.

This constellation of parameter alterations resolves the apparent paradox of why patients with an ostensibly cautious valuation profile engage in maladaptive risk-taking. With more concave utility for gains and more convex utility for losses, the Patient Group was, in terms of pure value computation, less sensitive to extreme outcomes; large potential rewards did not motivate them as strongly, and large potential losses did not deter them as effectively as they did for controls. However, their near-linear probability weighting meant they evaluated chance more objectively, without the adaptive pessimistic biases that typically protect against poor gambles. This combination leads to counterintuitive choices. For instance, a gamble with a small chance of a large gain but a high chance of a large loss would likely be rejected by a healthy individual, who would both subjectively overweight the high probability of the loss and feel its potential impact acutely. A patient with a frontal lesion, in contrast, might accept this gamble because they treat the probabilities more veridically and do not fully register the affective severity of the potential loss. Consequently, their risk-seeking behavior emerges not from a preference for risk itself, but as a by-product of attenuated sensitivity to outcome magnitudes coupled with a more literal processing of probabilities. They are not “risk lovers”; rather, their neural injury impairs their ability to value extreme outcomes and to deploy the cautious probability heuristics that support adaptive choice.

This interpretation aligns with and extends our understanding of frontal lobe function. The ventromedial prefrontal cortex (vmPFC) and orbitofrontal cortex (OFC) are critical for encoding the subjective value of options and integrating emotional signals into choice (Hiser & Koenigs, 2018; Messimeris et al., 2023). Damage to these areas likely underpins the abnormal utility curvature we observed; indeed, prior lesion studies have demonstrated that vmPFC damage can alter risk preferences and value computation (Pujara et al., 2015). The trend toward reduced loss aversion is also consistent with the hypothesis that frontal damage blunts emotional responses to negative outcomes. In healthy individuals, physiological arousal, mediated by fronto-limbic circuits, contributes to robust loss aversion (Sokol-Hessner et al., 2013); patients may lack this affective “punch.” The normalization of the probability weighting function suggests that frontal lesions disrupt the network responsible for generating these adaptive distortions, which may involve regions such as the dorsolateral PFC and anterior insula (Rolls et al., 2022; Panidi et al., 2022). By losing this regulatory frontal input, patients may default to a more mathematically “rational” but behaviorally maladaptive appraisal of chance. Intriguingly, this echoes findings that individuals with certain brain lesions can make more “objective,” expected-value-driven decisions in laboratory settings (Shiv et al., 2005). Our results demonstrate that this spurious rationality, when combined with an impaired appreciation of consequences, creates a uniquely hazardous decision-making profile.

Beyond these aggregate group differences, our study revealed a striking behavioral dissociation in how patients behaved across social contexts. They exhibited greater risk-taking when choosing for themselves but became more risk-averse when choosing for another person. This divergence is particularly notable because our control participants showed no such self-other gap, a finding consistent with some studies in healthy populations (Polman & Wu, 2020; Fareri et al., 2022). Importantly, this behavioral bifurcation does not stem from a context-dependent computational failure. Our Prospect Theory modeling revealed no significant Group × Context interactions on the core parameters (α, β, γ, λ), indicating that the underlying computational disruptions (specifically the blunted sensitivity to outcome magnitudes with lower α and β values, and more linear probability weighting with higher γ) were uniformly present in both self and other contexts. Rather, the behavioral dissociation suggests that this stable computational deficit interacts with context-specific motivational factors. The natural human tendency toward caution and responsibility when making decisions for others may have partially masked the computational deficit in the “Other” condition, leading to conservative rejection of most gambles. In contrast, the same computational deficit produced overt maladaptive risk-taking in the “Self” condition, where no such protective tendency exists and where even the diminished motivational pull of potential gains was sufficient to drive acceptance of poor gambles. Notably, our control participants showed stable decision-making across both contexts in terms of both computational parameters and behavior, confirming that healthy frontal function maintains consistent valuation processes regardless of social context. The fact that the patients’ disrupted computational system produced context-dependent behavioral outputs indicates a loss of the ability to flexibly adapt decision strategies while maintaining stable underlying valuations, a hallmark of frontal lobe dysfunction. another explanation for this self-other reversal is that decisions for another person engaged a more deliberative, social rule-based cognitive strategy that partially compensated for the patients’ emotional and valuation deficits. A patient might consciously invoke a social norm, such as a “do no harm” principle, and thus systematically reject risky options when acting on behalf of another, even while intuitively or automatically accepting similar gambles for themselves based on their disrupted valuation system. This aligns with the concept of responsibility aversion, where individuals become more cautious when their choices affect others’ outcomes (Edelson et al., 2018). Despite their injury, our patients may retain an intellectual understanding of this responsibility, and in the absence of the personal lure of a reward, they may default to an overly cautious strategy.

A complementary explanation lies in the differential affective engagement between contexts. In the “Self” condition, patients may still experience a diminished but present motivational pull from the possibility of winning. In the “Other” condition, this vicarious reward is likely even weaker. Lacking both a strong fear of loss and a personal excitement for gain, patients may simply find gambles for another person “not worth it,” leading to a conservative pattern of rejection. This interpretation is supported by our finding of reduced sensitivity to gain magnitude in patients; if large gains are not strongly motivating for oneself, they are likely even less so when they are for an anonymous other. Superficially, this pattern might suggest that patients are more altruistic or prosocial than expected, as they appear to exercise greater caution when another person’s welfare is at stake. However, our computational modeling reveals that this is not genuine altruism driven by empathic concern or other-regarding preferences. Rather, it reflects a computational artifact: the combination of blunted affective engagement with vicarious rewards and the preserved capacity to invoke social rules (e.g., “do no harm”) produces a behavioral pattern that mimics altruism without the underlying prosocial motivation. True altruism would involve a positive valuation of others’ gains; here, patients simply find others’ outcomes insufficiently motivating to justify risk, leading to blanket rejection rather than thoughtful prosocial choice.

Our results extend previous research by demonstrating that frontal lobe damage produces a context-dependent dysregulation of risk-taking, rather than a monolithic increase in impulsivity. Prior lesion studies have variously reported that frontal patients can be risk-seeking, apathetic, or show impaired social judgment (Koenigs & Tranel, 2007; Skromanis et al., 2023). By examining decisions in both personal and interpersonal contexts, we help reconcile these observations. Frontal patients can be both risk-seeking and risk-averse: they take excessive risks when only their own welfare is at stake, but become highly guarded when responsible for another’s outcome. This behavioral bifurcation underscores the critical role of the frontal lobes in supporting context-dependent cognitive flexibility.

These findings have significant practical implications. Individuals with frontal damage are likely prone to making poor personal financial choices, as they may underestimate the danger of high-risk propositions. Clinicians and caregivers should be aware that this behavior stems from an altered internal valuation system, not merely ignorance. Conversely, the same patient might become excessively cautious or indecisive when making decisions for others, potentially forgoing beneficial opportunities. Rehabilitation should therefore aim to restore balance by encouraging more caution in personal decisions and appropriately calibrated confidence in social ones.

Our results suggest several targets for cognitive rehabilitation. Metacognitive strategy training could teach patients to pause and explicitly analyze high-stakes choices, compensating for their impaired intuitive guidance. Given their more linear probability weighting, patients may benefit from being explicitly trained to consider worst-case scenarios, a cognitive step they may no longer perform automatically. Furthermore, emerging evidence suggests that noninvasive brain stimulation of prefrontal regions can modulate decision biases (Panidi et al., 2022; Kroker et al., 2023), offering a potential future avenue for therapeutic intervention.

This study has limitations. The patient sample was heterogeneous, and the sample size was moderate. Future work using voxel-based lesion-symptom mapping in larger cohorts could test whether specific parameter changes map onto distinct frontal subregions. For instance, damage to vmPFC may drive the observed changes in utility curvature, while damage to dorsomedial PFC, a region involved in social cognition (Piva et al., 2019), might specifically contribute to the abnormal self-other dissociation. Additionally, future studies could incorporate more complex paradigms to examine learning and feedback processing or explore decisions in more naturalistic social contexts.

In conclusion, this study demonstrates that frontal lobe lesions produce a specific and consequential alteration in the computational architecture of risky choice. Patients with frontal damage exhibit a unique profile of blunted sensitivity to outcome magnitudes combined with an unusually linear perception of probabilities. This profile explains their paradoxical tendency to make risky choices for themselves while becoming overly cautious for others. By providing a precise, mechanistic account of these deficits, our findings deepen the understanding of the frontal lobes’ contribution to both individual and social decision-making and point toward targeted interventions that could improve the quality of life for individuals with brain injury.

